# Myeloperoxidase enhances the migration of human choriocarcinoma JEG-3 cells^1^

**DOI:** 10.1101/2023.01.28.526039

**Authors:** ZN. Mihalic, T. Kloimböck, N. Cosic-Mujkanovic, P. Valadez-Cosmes, K. Maitz, O. Kindler, C. Wadsack, A. Heinemann, G. Marsche, M. Gauster, J. Pollheimer, J. Kargl

## Abstract

Myeloperoxidase (MPO) is one of the most abundant proteins in neutrophil granules. It catalyzes the production of reactive oxygen species, which are important in inflammation and immune defense. MPO also binds to several proteins, lipids, and DNA to alter their function. MPO is present at the feto-maternal interface during pregnancy, where neutrophils are abundant. In this study, we determined the effect of MPO on JEG-3 human choriocarcinoma cells as a model of extravillous trophoblasts (EVTs) during early pregnancy. We found that MPO was internalized by JEG-3 cells and localized to the cytoplasm and nuclei. MPO internalization and activity enhanced JEG-3 cell migration, whereas this effect was impaired by pre-treating cells with heparin, to block cellular uptake, and MPO-activity inhibitor 4-ABAH. This study identifies a novel mechanism for the effect of MPO on EVT function during normal pregnancy and suggests a potential role of MPO in abnormal pregnancies.

**GRAPHICAL ABSTRACT:** 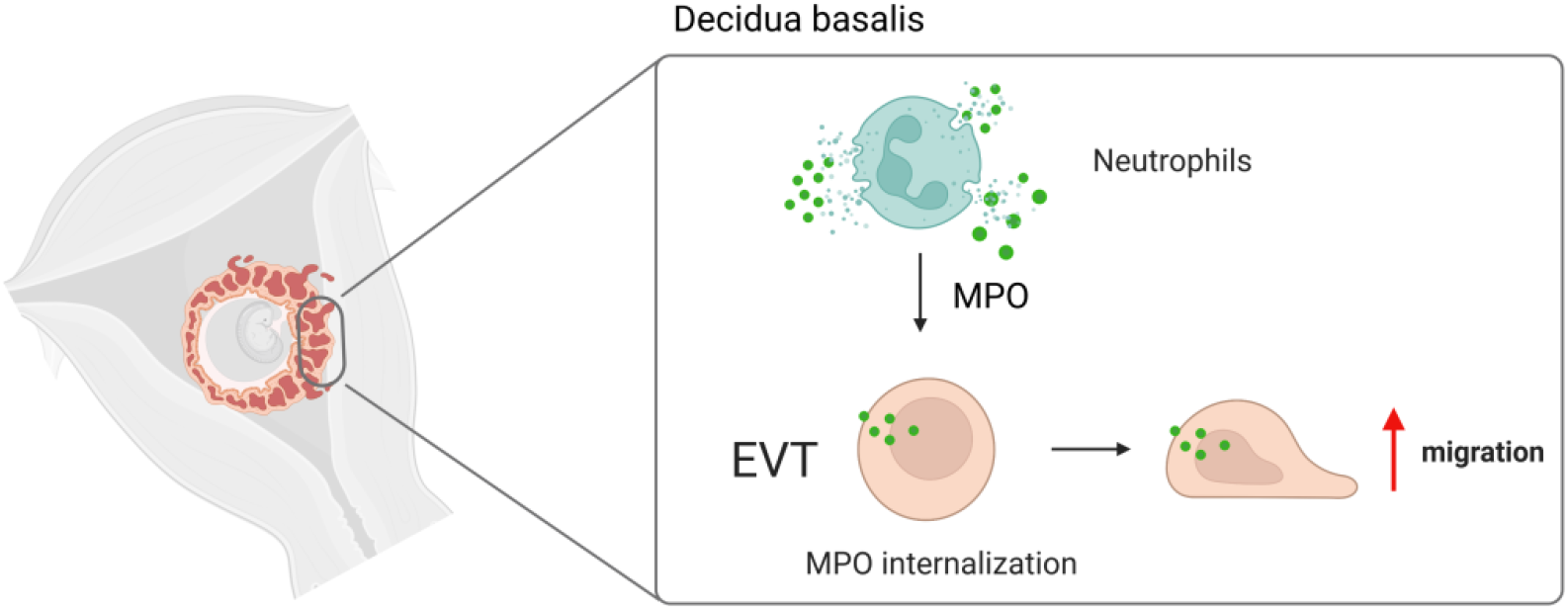

## INTRODUCTION

Pregnancy is an immunological challenge, in which the mother and fetus must coexist. During early pregnancy, placenta-derived extravillous trophoblasts (EVTs) invade the decidua, engraft and remodel uterine spiral arteries. They gradually replace maternal endothelial cells to enable an adequate blood flow into the placenta (1). Therefore, it is important that this process is highly regulated as defective EVT invasion results in abnormal placentation. Maternal immune cells represent 30%–40% of all decidual cells (2). Maternal leukocytes, particularly uterine natural killer cells, regulate EVT invasion and are important for normal development during pregnancy (3). However, other maternal immune cells, such as neutrophils, are also required for the development of a healthy feto-maternal interface, including neutrophils, which are highly present in the decidua basalis (DB) (4). Neutrophils are part of the innate immune system and are one of the first cell types that respond to acute inflammation. The infiltration of neutrophils during pregnancy occurs at the decidua basalis (DB) (5,6). Depletion of neutrophils in mouse models results in abnormal placentation (7); however, the role and interactions of neutrophils with EVTs are still unclear.

Myeloperoxidase (MPO) is a heme-bound enzyme primarily expressed in the azurophilic granules of neutrophils, but also present in monocytes and macrophages (8). It represents 5% of the whole neutrophil cell weight, making it one of the most abundant proteins in neutrophils (9). Upon neutrophil activation, MPO is transported into phagosomes or released into the extracellular space (10). It catalyzes the production of reactive oxygen species (ROS), such as hypochlorous acid, in an oxidative reaction of hydrogen peroxide with halides (chloride, bromide, iodide), which are important for pathogen killing during infection (11). MPO is highly positively charged because of its high arginine and lysine content; therefore, it also binds to a variety of proteins, DNA, and lipids (12). Besides its role in pathogen killing, MPO has been shown to affect neutrophil recruitment by binding to CD11b/CD18 integrins (13), proteins, and DNA, as well as to play a role in lipid modification, regulation of apoptosis, neutrophil extracellular trap (NET) formation, and interactions with macrophages (8). Furthermore, MPO affects angiogenesis of endothelial cells (14).

In the placenta, the presence of neutrophils and MPO in the DB has been confirmed (4,5,15). Furthermore, higher MPO levels on the surface of blood neutrophils have been observed during pregnancy (16); however, its abundance and role during first trimester implantation and placentation remains unclear. Therefore, we examined the role and uptake of MPO using the JEG-3 EVT model cell line. We hypothesize that MPO affects JEG-3 function in vitro and that its internalization by these cells is important for its effects. Our results indicate that JEG-3 uptake of MPO stimulates their migration, which may represent a drug target for pregnancy complications, associated with shallow trophoblast invasion.

## MATERIAL AND METHODS

### Cell culture

Human placental–epithelial choriocarcinoma cells (JEG-3 HTB-36TM ATCC®) were cultured in Earle’s Minimum Essential Medium (EMEM) (Gibco™) containing 2-mM L-glutamine supplemented with 1% nonessential amino acids (Gibco™), 1 mM sodium-pyruvate (GibcoTM), 10% fetal bovine serum (FBS) (Gibco™) and 1% penicillin/streptomycin (P/S) (Gibco™) at 37°C and 5% CO_2_. Before treatment, the cells were starved for 4 h in starvation medium (EMEM without FBS, P/S, nonessential amino acids, and sodium-pyruvate) and treated with the indicated concentrations of MPO (Elastin products company), in combination with 150 μg/ml heparin (Gilvasan, 1000 IE/ml) or 40-μM 4-aminobenzoic acid hydrazide (4-ABAH) (Sigma-Aldrich) where applicable.

### Immunofluorescence Staining

Paraffin-embedded placenta sections prepared from first trimester elective pregnancy terminations were heated at 60°C, immersed in xylene, rehydrated in descending alcohol concentrations (100%, 96%, 80%, 70%, 50%), washed in PBS without Ca^2+^ and Mg^2+^ (PBS−/−), and boiled in antigen retrieval buffer (2.94-g/L tri-Natrium-Dihydrat, MW 294 g, pH=6) for 15 min. Cold antigen retrieval buffer was added and cooled to 4°C for 1 h. The slides were washed in PBS−/−, incubated for 30 min in 0.3% H_2_ O_2_ (in PBS−/−), and subsequently washed in PBS−/−. The slides were incubated in blocking solution [0.3% TRTX, 5% goat serum, 4% BSA (Sigma-Aldrich) in PBS−/−] at room temperature for 1.5 h and subsequently incubated with primary rabbit anti-MPO antibody (Cell Signaling, clone: E1E7I, 1:500) and primary mouse anti-HLA-G antibody (BD, clone: 4H84, 1:2500) in a humidified box at 4°C overnight. The slides were washed again and incubated in goat antirabbit AlexaFluor488 (Invitrogen, 1:500) and goat anti-mouse AF647 (Invitrogen, 1:500) secondary antibody for 3 h in the dark. The slides were then washed in PBS−/− and background was reduced using Vector True View Kit mix for 3 min. The slides were washed in PBS−/− and the nuclei were stained with Vecta Shield Mounting Medium containing DAPI (Szabo Scandic).

### MPO RNA expression

Basal plate RNA expression data (17) was acquired from the dataset GSE22490 of the NCBI Gene Expression Omnibus database (GEO) (http://www.ncbi.nlm.nih.gov/geo/) via the R package GEOQuery (n=10). MPO expression was plotted with the help of R packge ggplot. (R version 4.2.2) (18,19).

### Immunocytochemistry

JEG-3 cells were seeded into 6-well plates on coverslips in growth medium, incubated for 24 h, and subsequently starved overnight. Afterwards, the cells were treated in starvation medium with 10 μg/ml of MPO (Elastin Products Company) or 5 μg/ml MPO in combination with heparin and 4-ABAH. Heparin and 4-ABAH were added directly prior to MPO treatment and incubated for 4 h. Afterwards, the coverslips were washed with PBS−/−, the cells were fixed with 100% ice-cold methanol for 20 min at − 20°C, and subsequently washed with PBS−/−. Blocking was performed in PBS−/− supplemented with 5% goat serum (Sigma-Aldrich) and 5% BSA (Sigma-Aldrich) for 1 h at RT. A primary rabbit anti-MPO antibody (Cell Signaling, 1:500) in 1:10 blocking solution + 0.1% Triton X-100 was added and incubated overnight at 4°C. After washing with PBS−/−, the coverslips were incubated with secondary antibody (goat antirabbit AlexaFluor488 (Invitrogen, 1:500) for 1 h at RT followed by another washing step. The coverslips were mounted along with DAPI (Vecta Shield Mounting Medium) on glass slides. The cells were observed using an Olympus IX70 fluorescence microscope equipped with a Hamamatsu ORCA-ER digital camera (Hamamatsu Photonics K.K., Japan). CellSense 1.17 Dimension software was used for processing the images.

### Bromodeoxyuridine proliferation assay

Following treatment with MPO [V (EMEM -/-), 0.5, 2, 5 µg/ml] for 24, 48, and 72 h, 10-μM bromodeoxyuridine (BrdU) solution was added 4 h before the end of the experiment [fluorescein isothiocyanate (FITC) BrdU Flow Kit (BD #559619)] according to the manufacturer’s protocol. Briefly, the cells were washed in PBS−/− and stained with Fixable Viability Dye eFluor™ 450 (Thermo Scientific) for 20 min. Afterwards, the cells were washed in PBS−/− and permeabilized using Cytofix/Cytoperm buffer (BD) for 20 min. Following fixation, the cells were washed in 1X BD Perm/Wash Buffer and treated with 30-μg/ml DNAse I for 1 h at 37°C. The cells were washed and stained with FITC-conjugated BrdU Antibody (1:50) prepared in Perm/Wash buffer (BD) for 20 min. Upon stimulation, the cells were again washed, and proliferation was measured by flow cytometry.

### Annexin V apoptosis assay

After treatment with 5 µg/ml MPO for 6, 15, 18, and 24 h, the cells were stained in 1x Annexin V Binding Buffer and incubated with FITC-anti human Annexin V antibody (1:50) and PI (1:50) for 15 min, according to manufacturer’s instructions as previously described (20).

### Scratch assay

Cells (6 × 10^5^/well) were seeded into 6-well plates, grown for 24 h, and subsequently starved in starvation medium for 5 h. The starved cells were washed with PBS−/− and subsequently scratched using a SPLScarTM Scratcher suitable for 6-well plates (SPL Life Science). The cells were washed and treated with 0 μg/ml, 2.5 μg/ml or 5 μg/ml MPO. Light microscopy images were captured at 100x magnification (Nikon) directly after wounding and after 24 h. To calculate the amount of wound closure, the cell-free area was determined using ImageJ software. Wound closure after 24 h is presented as the fold-change percentage of the initial wound area.

### Electric cell-substrate impedance sensing migration assay

Electric cell-substrate impedance sensing (ECIS) was performed in a 96W1E+ plate (96-well array from Applied BioPhysics). The wells were activated using 10-mM L-cysteine (Sigma-Aldrich) for 10 min at RT. Afterward, the wells were washed twice with water and once with growth medium. Cells (4.8 × 10^5^/well) were seeded and incubated for 48 h. Confluent cells were starved overnight. Resistance was measured at 4000 Hz (ECIS® Ζθ) and wounding was performed in each well at 3,000 μA and 48,000 Hz for 30 sec. Afterwards, the cells were treated with 0 μg/ml, 1 μg/ml, 2.5 μg/ml, 5 μg/ml, and 10 μg/ml MPO, and 5 μg/ml MPO in combination with heparin and 4-ABAH. Resistance was measured for 24 h in 60 s intervals and expressed as fold-change relative to the baseline resistance before wounding. Post-wounding values were subtracted for normalization and represented as mean of at least eight replicates per condition in each individual experiment.

### Nuclear and cytoplasmic fractionation of JEG-3 cells

Cells (5 × 10^5^) were seeded into 10 cm dishes and incubated until confluent. The cells were starved for 4 h and subsequently incubated with 10 µg/ml MPO for 2 h. Afterwards, the cells were washed in PBS−/−, 1 ml of ice-cold PBS−/− was added, and the cells were scraped. The cells were pelleted for 5 min at 13,000 rpm (4°C) and the supernatants were aspirated. Next, 500 µl Buffer A [10 mM HEPES, 10 mM KCl, 0.1 mM EDTA, protease/phosphatase inhibitor (100×), (Cell Signaling) in PBS−/− was added. Cells were homogenized for 15 s at medium power on ice. Then, 25-μl NP-40 was added and thoroughly mixed by vortexing. Lysates were centrifuged for 5 min at 5000 rpm at 4°C. Supernatants were separated (cytoplasmic fraction) from the remainder and stored on ice. For nuclear fractions, the pellets were mixed with 50 µl ice-cold buffer C (20-mM HEPES pH 7.9, 25% glycerol, 0.4-M NaCl, 1-mM EDTA in PBS -/-). The pellet was gently dislodged and afterwards vigorously rocked in ice for 30 min. Lysates were centrifuged for 10 min at 14,000 rpm at 4°C. The supernatant (nuclear fraction) was collected and both fractions were used for western blot analysis.

### Western blot analysis

Total protein was isolated from frozen cells in IP buffer (0.1% Triton X-100, 150-mM NaCl, 25-mM KCl, 10-mM Tris HCl, pH 7.4; 1-mM CaCL_2_ in H_2_ O) supplemented with 1:100 protease/phosphatase inhibitor cocktail (Cell Signaling). The protein content was measured using a Pierce BCA assay kit (Thermo Scientific). The samples were mixed with 4× NuPage sample buffer (90-μl 4× LDS sample buffer + 10-µl reducing reagent) and boiled for 10 min at 95°C. Next, 20 µg of protein from each homogenate was applied to the NuPage, Bis-Tris gel (Invitrogen) and electrophoresed for 45 min at 200 V. The proteins were transferred to a membrane using an iBlot transfer device and blocked in TBST (1× TBS + 0.1% Tween) supplemented with 5% milk for 1 h. The membrane was incubated with the primary antibody overnight at 4°C. Anti-MPO (rabbit antihuman MPO, clone: E1E7I, 1:500 dilution, Cell Signaling), anti-GAPDH (rabbit antihuman GAPDH, clone: 14C10, 1:500 dilution, Cell Signaling), anti-Lamin A/C (mouse antihuman Lamin A/C, clone: 4C11, 1:1000 dilution, Cell Signaling), and anti-tAKT (rabbit antihuman tAktin, clone: 11E7., 1:1000 dilution, Cell Signaling) monoclonal antibodies were used. The membranes were washed for 30 min and subsequently incubated with secondary antibody HRP-Goat anti-rabbit IgG (polyclonal, 1:5000 dilutions, Jackson ImmunoResearch) for 2 h at RT. The membranes were washed and developed using ECL solution (BioRad).

### Flow cytometry

Cells (3 × 10^5^/well) were seeded into 6-well plates and incubated for 24 h. The cells were then starved overnight and treated with 2.5- and 5-μg/ml MPO or MPO in combination with heparin and 4-ABAH for 4 h. The cells were fixed and permeabilized for 40 min using a BD transcription factor fixation/permeabilization kit to allow the antibodies to bind to nuclear protein. Subsequently, the cells were preincubated with Fc receptor blocking solution followed by antihuman MPO-FITC-conjugated antibody (DB) for 30 min at 4°C, washed twice in TF washing buffer (BD), and once in PBS without PBS−/− supplemented with 2% FBS. Samples were measured using a Canto flow cytometer with FACSDiva software (BD). Analysis and compensation were performed using FlowJo software (TreeStar).

### Myeloperoxidase activity

MPO activity was measured as previously described (21). Briefly, cells (3.5 × 10^5^/well) were seeded into 6-well plates in 2ml EMEM+/+ and incubated for 24 h. The cells were then starved for 4 h and treated for 4 h with 5 µg/ml MPO. The cells were detached with accutase (Sigma-Aldrich) and washed in PBS−/−. The cells were resuspended in 500 µl inhibitor cocktail (1:100 dilution in PBS−/−), sonicated for 15 s on ice and centrifuged for 10 min at 18,000 rpm (4°C). In a 96-well plate, 100 µl of 3,3′,5,5′-tetramethylbenzidine solution (Thermo Fisher), 2 µl of H_2_ O_2_ (1:300 of 30% H_2_ O_2_), and 5 µl of sample supernatant were added. The reaction was incubated for 30 min in the dark and stopped with 100 µl 2-N sulfuric acid. The absorbance was measured at 450nm.

## RESULTS

### Neutrophils are located near extravillous trophoblasts in first trimester decidua basalis

There is little information regarding the localization and interaction of neutrophils and EVTs in the DB during the first trimester of pregnancy. Therefore, we determined the localization of MPO, used as a marker for neutrophils, and EVTs (detected by HLA-G) in first trimester DB. We observed MPO-positive neutrophils (green) in close proximity to HLA-G positive EVTs (magenta) via fluorescence immunohistochemistry (Figure 1a). Consistent with our data, it has been previously shown that MPO accumulates in human placenta and the DB in the third trimester of pregnancy (15). Further, we investigated a publicly available bulk RNAseq data set for the expression of MPO in first trimester basal plate (17). We observed MPO expression in all samples (Figure 1b). Each bar represents a separate sample.

**Figure 1.**
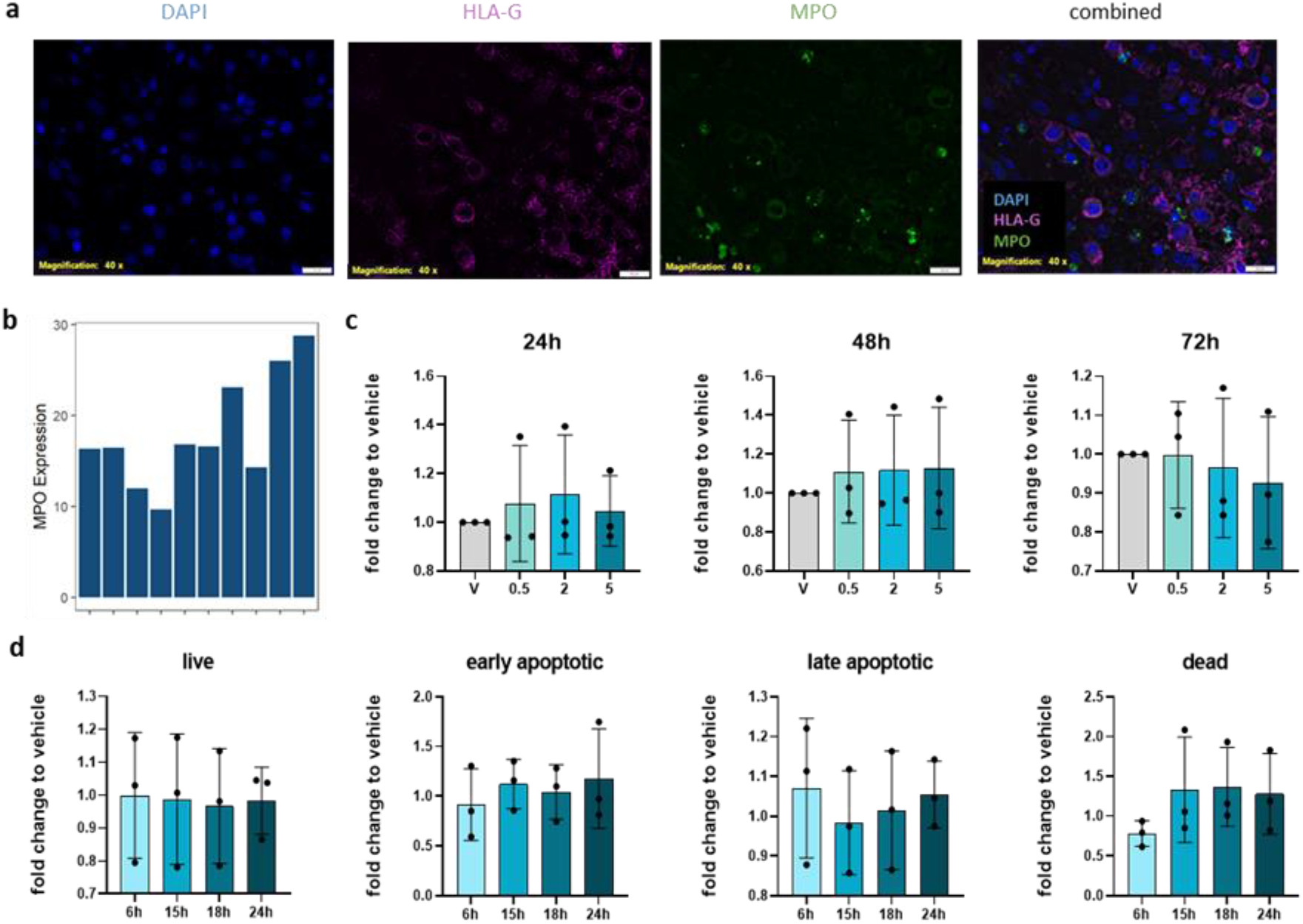
MPO is present in decidua basalis. **a)** Representative immunofluorescence microscopy of first trimester decidua basalis. Slides were stained for HLA-G (magenta) and MPO (green). b) MPO expression in basal plate of publicly available RNASeq data. Each bar represents a separate sample (N=10). c) Proliferation of JEG-3 cells after 24 h, 48 h, and 72 h treatment with vehicle or 0.5 µg/ml, 2 µg/ml, and 5 µg/ml MPO (N=3). d) JEG-3 Annexin V/PI apoptosis assay. Live, early apoptotic, late apoptotic, and dead cells were measured by flow cytometry following treatment with 5 µg/ml MPO. Annexin V and PI were measured at 6 h, 15 h, 18 h, and 24 h following MPO treatment and compared with vehicle-treated cells (N=3). One-way analysis of variance (ANOVA) and Tukey’s post hoc test were performed. Data are represented as the mean ± SD, *P < 0.05.

### Myeloperoxidase increase JEG-3 migration in vitro

The JEG-3 choriocarcinoma cell line serves as a model to study EVT cell behavior in the presence of MPO as primary trophoblasts lack sufficient proliferation in vitro (22). To assess the effect of MPO on JEG-3 cells, we exposed them to various concentrations of MPO and measured the effect on cell proliferation after 24 h, 48 h, and 72 h of treatment. We did not observe any differences in proliferation of JEG-3 cells comparing vehicle to increasing concentrations of MPO (0.5, 2, and 5 μg/ml) at any timepoint (Figure 1c). Furthermore, we performed AnnexinV/PI staining to determine whether MPO affects JEG-3 apoptosis. We measured live (AnnexinV− /PI−), early apoptotic (AnnexinV+/PI−), late apoptotic (AnnexinV+/PI+), and dead (AnnexinV− /PI+) cells at 6 h, 15 h, 18 h, and 24 h of treatment (Figure 1d). Similar to the proliferation assay, we did not observe any effect of MPO on JEG-3 apoptosis compared with untreated cells.

During normal/healthy early pregnancy, EVTs migrate and invade in large numbers toward the maternal spiral arteries to supply adequate blood flow to the fetus (23). During pregnancy complications, such as pre-eclampsia and intrauterine growth restriction (IUGR), EVTs do not invade deeply enough, which affects spiral artery remodeling (24). Because EVT migration is a key feature in normal placentation, we determined whether MPO affects this process. First, we conducted a wound-healing assay using JEG-3 cells (Figure 2a). JEG-3 cells treated with 2.5 μg/ml MPO did not significantly increase wound closure after 24 h; however, there was a trending increase in migration of treated cells compared with the control. Upon treatment with 5 μg/ml MPO, JEG-3 cells showed significantly enhanced levels of migration compared with the untreated group (Figure 2b). To confirm this effect, we performed an ECIS migration assay at four different MPO concentrations (1–10 μg/ml) for up to 24 h (Figure 2c). At 14 h post-wounding, enhanced wound closure was observed in MPO-treated JEG-3 cells compared with the control cells (Figure 2d).

**Figure 2.**
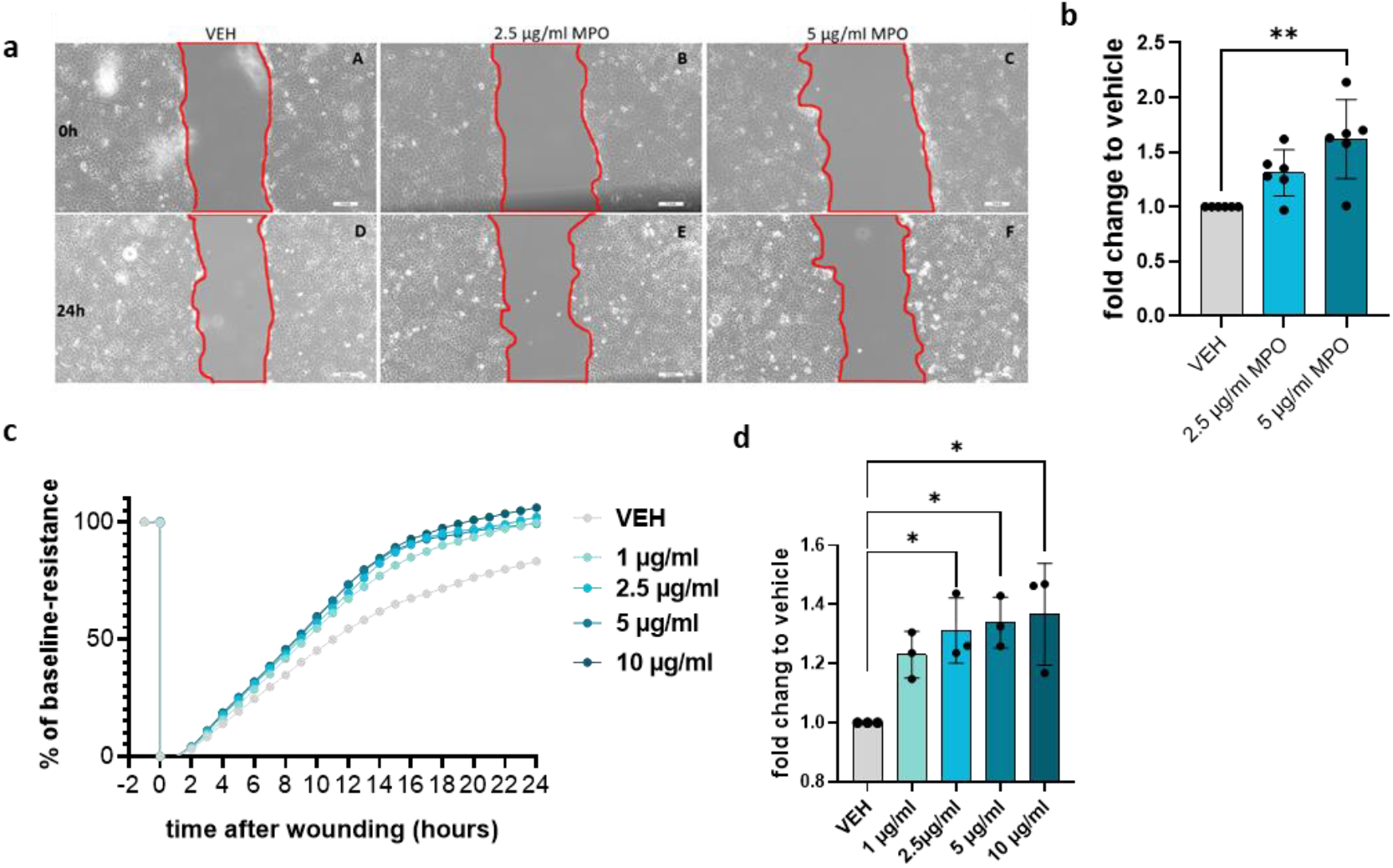
MPO alters JEG-3 migration in vitro. **a)** Scratch migration assay of JEG-3 cells treated with vehicle, 2.5 µg/ml, and 5 µg/ml MPO. Microscopic images were taken immediately (A–C) and 24 h (D–F) after wounding. **b)** Scratch assay results are presented as % cell-free area at time 0 and after 24 h normalized to the vehicle-treated cells (N=6). **c)** Wound closure capacity of JEG-3 cells measured by ECIS (N=3) after treatment with 1 µg/ml, 2.5 µg/ml, 5 µg/ml, and 10 µg/ml MPO. The cell monolayer was wounded at 3000 µA and 48,000 Hz for 30 sec (t = 0). Post-wounding resistance values were normalized to the wounding efficiency by subtracting the resistance value after wound induction from all values. Wound closure was monitored over 24 h and expressed as fold-change compared with baseline resistance before wounding. (baseline resistance: 1 = monolayer, 0 = wounding). **d)** Results of ECIS measurements represented as mean fold-change relative to the vehicle after 14 h. One-way ANOVA and Tukey’s post hoc test were performed. Data are presented as the mean ± SD, *P < 0.05.

### Myeloperoxidase uptake by JEG-3 cells and nuclear localization

MPO binds to the surface of endothelial and epithelial cells through binding to glycosaminoglycans and is rapidly internalized by the exposed cells (25). To determine whether MPO is taken up by JEG-3 cells, we treated JEG-3 cells with increasing MPO concentrations, subsequently cells were washed thoroughly to remove remaining MPO in the media and measured internalized MPO protein levels via western blot analysis. Total MPO protein amount increased in JEG-3 cells treated with higher concentrations (Figure 3a). Immunofluorescence microscopy of vehicle and MPO-treated JEG-3 cells revealed that untreated cells did not contain MPO. Upon treatment, MPO was internalized and localized to the cytoplasm and nuclei (Figure 3b). Furthermore, we observed that not all the cells bind or take up MPO, which was confirmed by flow cytometry (Figure 3c and d). We stained for MPO following nuclear permeabilization and observed that upon treatment with 2.5 µg/ml and 5 µg/ml of MPO, approximately 4% and 9% of the cells, respectively, internalized MPO (Figure 3c and d). To assess MPO intracellular localization following uptake, the cells were separated into cytoplasmic and nuclear fractions following 5 µg/ml MPO treatment for 2 and 24 h. MPO was observed in both the cytoplasmic and nuclear fraction at each time point (Figure 3e). To confirm the purity of these fractions, Lamin A/C was present only in the nuclear fraction and tAKT was more abundant in the cytoplasmic fraction (Figure 3e). MPO activity assays revealed that MPO was active upon cell uptake, measured in cell lysates after 4 h of treatment (Figure 3f).

**Figure 3.**
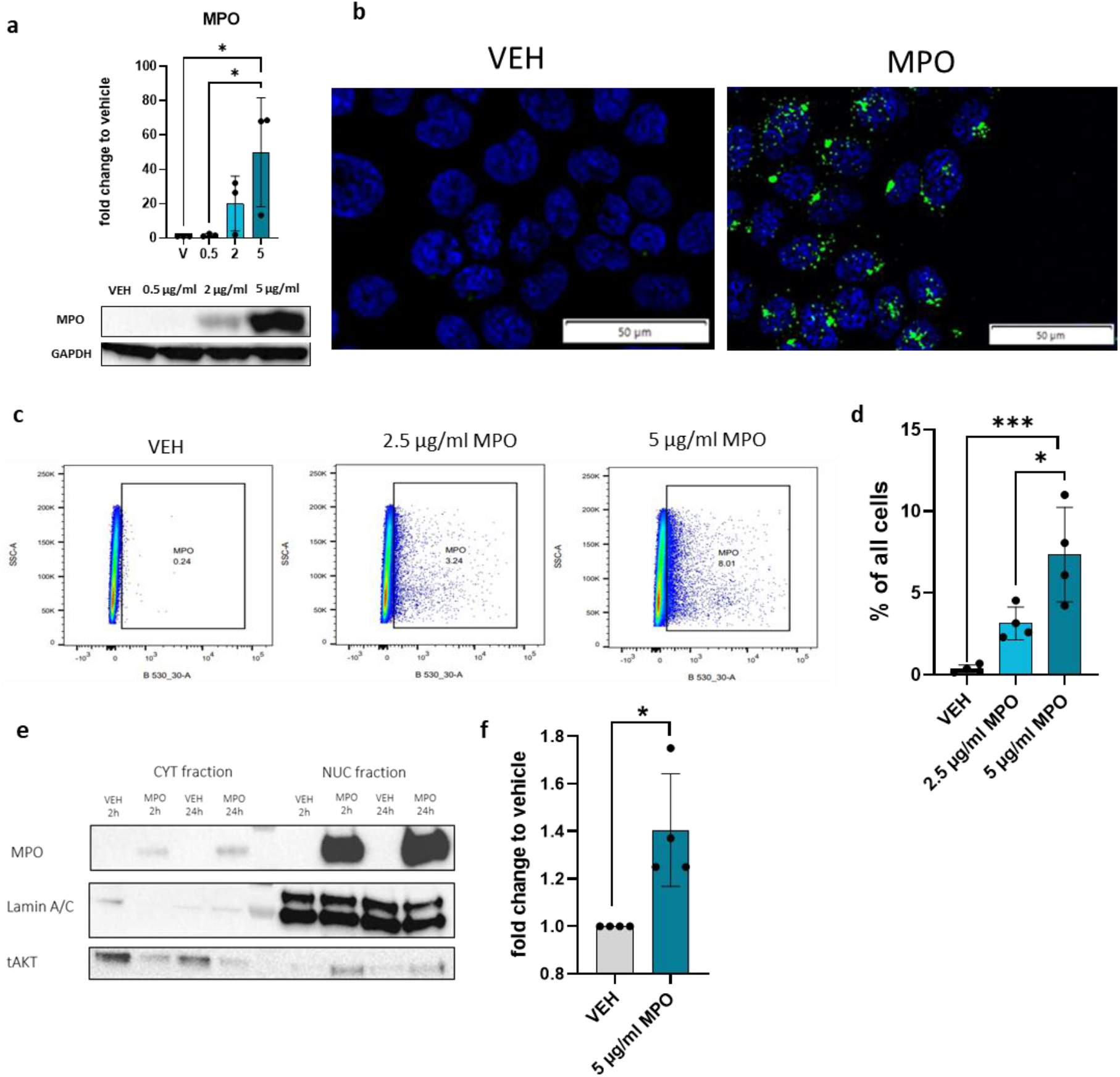
MPO uptake by JEG-3 cells and nuclear localization. **a)** Western blot analysis of JEG-3 cells treated with various concentrations of MPO (vehicle, 0.5 µg/ml, 2 µg/ml and 5 µg/ml) and presented as fold-change relative to the vehicle (N=3). **b)** Representative immunofluorescence microscopy photos of vehicle and 10µg/ml MPO-treated cells. The cell nucleus was stained with DAPI and MPO is represented in green with AF488-labeled secondary antibody. **c)** Scatterplots representing vehicle control, 2.5 μg/ml MPO, and 5 μg/ml MPO-treated JEG-3 cells. MPO-positive cells were gated based on the vehicle. Cells were treated for 4 h and intranuclear-stained for flow cytometry using an FITC-anti-MPO antibody (N=4). **d)** Results of flow cytometry analysis represented as % of all cells. e) Representative western blot of JEG-3 cell fractions stained for MPO, Lamin A/C, and tAKT of vehicle and 5 μg/ml MPO-treated cells. Cytoplasmic and nuclear fractions were isolated. f) MPO activity assay of MPO-treated JEG-3 cell lysates presented as fold-change relative to the vehicle-treated cells (N=4). One-way ANOVA and Tukey’s post hoc test were performed for multiple comparisons. To compare two samples, a Mann–Whitney paired t-test was performed. Data are presented as the mean ± SD, *P < 0.05.

To determine whether this effect was observed in other invasive trophoblast cell lines, we performed similar experiments on the ACH-3P human trophoblast cell line (26). Consistent with the JEG-3 results, an increase in total MPO in ACH-3P cells was observed by western blot analysis (Supplementary Figure 1a). Furthermore, immunofluorescence microscopy showed localization of MPO in the cytoplasm next to the nuclei (Supplementary Figure 1b). Upon fractionation, we observed increased MPO localization in treated cells compared with the vehicle control. MPO was found in both cytoplasmic and nuclear fractions, and efficient fractionation was further confirmed by i) increased GAPDH present in the cytoplasmic fraction, ii) Lamin A/C present in the nuclear fraction, and iii) increased tAKT present in the cytoplasmic fraction (Supplementary Figure 1c).

### Myeloperoxidase uptake and activity is required for migration

MPO uptake by cells can be blocked by heparin, a member of the glycosaminoglycan (GAG) family (27).Further, MPO activity can be impaired by aminobenzoic acid hydrazine (4-ABAH), a potent irreversible inhibitor of MPO (28). Therefore, we determined the effect of these inhibitors on the internalization and function of MPO during JEG-3 migration. We treated JEG-3 cells with MPO, heparin plus MPO, or 4-ABAH plus MPO. Immunofluorescence microscopy confirmed that heparin prevented MPO internalization into JEG-3 cells, while 4-ABAH did not affect the internalization of MPO, as expected (Figure 4a). To determine the effect of heparin and 4-ABAH on MPO uptake, we performed flow cytometry analyses, measuring MPO levels in treated cells (Figure 4b). Following addition of heparin, MPO uptake was completely abolished compared with cells treated with MPO alone. In contrast, MPO uptake was not affected by 4-ABAH in the JEG-3 cell line (Figure 4b). To confirm these findings at the protein level, we performed western blot analyses. Cells were treated with MPO alone as a positive control, MPO plus heparin, or MPO plus 4-ABAH. Additionally, we heat-inactivated MPO to destroy its enzymatic activity and three-dimensional structure to block GAG interactions as a negative control (Figure 4c). MPO was not detected in heparin-treated samples, where uptake was abolished, and in samples where MPO was denatured. Furthermore, 4-ABAH exhibited a small decrease in MPO internalization (Figure 4d).

**Figure 4.**
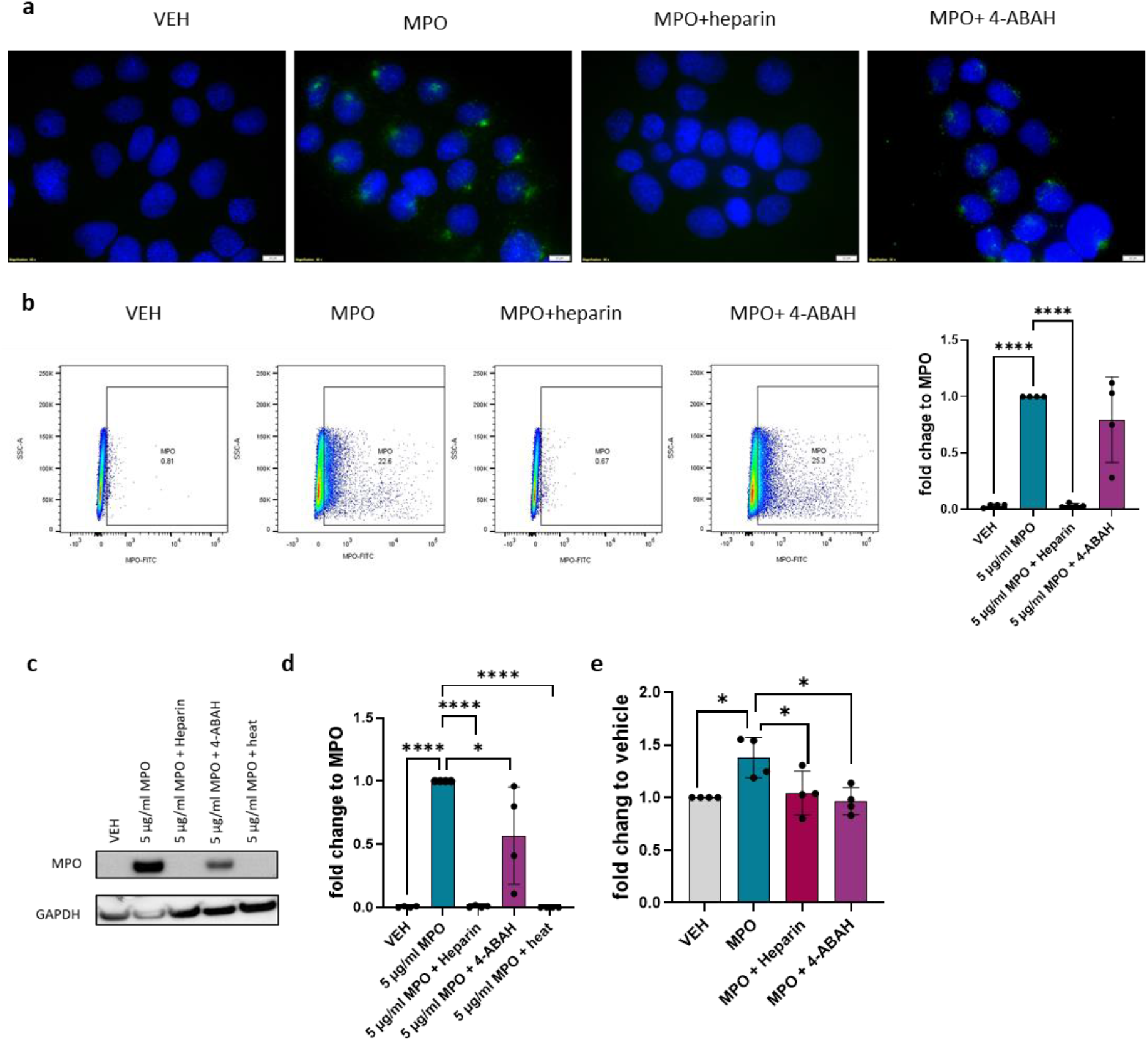
MPO uptake and activity are important for enhanced migration. **a)** Representative immunofluorescence photos of vehicle and 5 µg/ml MPO-treated cells in combination with heparin (blocks MPO uptake) and the MPO activity inhibitor 4-ABAH. The cell nucleus was stained with DAPI and MPO is represented in green, labeled with AF488 secondary antibody. **b)** Scatterplots representing vehicle control and 5 μg/ml MPO-treated JEG-3 cells in combination with heparin and 4-ABAH treatment. MPO-positive cells were gated based on vehicle-treated cells. Cells were treated for 4 h and intranuclear-stained for flow cytometry using an FITC-anti-MPO antibody (N=4). Flow cytometry analysis of MPO-positive cells presented as fold-change relative to MPO-treated cells (N=4). **c)** Representative western blot images of MPO and GAPDH of JEG-3 cells treated with vehicle and 5 µg/ml MPO in combination with heparin, 4-ABAH, or heat inactivated MPO. **d)** Western blot analysis presented as fold-change relative to MPO-treated cells (N=4). e) Results of ECIS measurements presented as mean fold-change relative to vehicle after 14 h (N=4). One-way ANOVA and Tukey’s test post hoc was performed for multiple comparisons. Data are presented as the mean ± SD, *P < 0.05.

Because we previously showed that MPO increases JEG-3 migration, we inhibited MPO uptake and activity using heparin and 4-ABAH. Most importantly, both heparin and 4-ABAH blocked MPO-induced cell migration (Figure 4e).

## DISCUSSION

Studies of neutrophils at the feto-maternal interface indicate that proper neutrophil infiltration has an important role in normal development of the fetus. In recent mouse studies, depletion of neutrophils using anti-Gr1 or anti-Ly6G antibodies resulted in inadequate blastocyst implantation, as well as altered placenta development in pregnant mice (7,29), suggesting that neutrophils are important at the early stages of pregnancy. In the present study, we determined whether MPO, predominantly expressed by neutrophils and released into the extracellular space, plays a role during placentation. The JEG-3 choriocarcinoma cell line was used as a model to study EVT cell behavior in the presence of MPO, as primary trophoblasts lack efficient proliferation in vitro.

First, we showed that MPO was present and in proximity to EVTs in the DB. MPO expression in the DB during early and late gestation has previously been described (15,30). Blood neutrophils from pregnant women accumulated higher amounts of MPO at the surface and released more ROS in comparison with that of nonpregnant women (16). In some cancers, MPO increases tumor progression and metastases resulting in a higher migration of tumor cells (31). In endothelial cells, MPO promotes tube formation and increases wound healing (14). Similarly, we demonstrated that JEG-3 cells treated with MPO exhibit a higher migratory ability compared with vehicle-treated control cells. In addition, proliferation and apoptosis were not influenced by MPO, which suggests that the migratory effect is due to MPO activity and not the direct result of proliferating cells. Similar to our data, MPO did not induce apoptosis in several endothelial cell lines; however, the same study found that MPO was internalized and was present in the cytoplasm and nucleus of endothelial cells and that there were higher levels of intracellular oxidants (32). We found that total MPO within the cells was increased at higher concentrations of MPO. During inflammation, levels of MPO are reported to be elevated (33), but there is no data, to our knowledge, regarding local MPO concentrations around neutrophils following degranulation. We hypothesize that upon activation of neutrophils, there is a high concentration of MPO released in their surrounding environment. In addition to degranulation and release of MPO and other granular proteins into the extracellular space, neutrophils can undergo NETosis, leading to accumulation of extracellular MPO bound to DNA (34). Consistently, MPO was taken up by JEG-3 cells and localized in the cytoplasm and the nucleus as determined by immunofluorescence and cell fractionation studies. The same effect was observed in the ACH-3P EVT cell line. These data indicate that MPO uptake is not a cell-specific effect, but rather a more general MPO effect. In contrast, previous reports of the effect of activated neutrophils on the Swan 71 trophoblast cell line showed a reduction in EVT migration. In this study, NET formation was induced, which suggests that MPO bound to DNA could not be internalized through glycosaminoglycan binding (35). The neutrophil inflammatory state may be a key event in understanding neutrophil-EVT interactions. Depending on their pro- or anti-inflammatory phenotype, neutrophils can release different molecules, which depend upon microenvironmental stimuli (36). During pregnancy, neutrophils promote Treg differentiation, favoring immune tolerance (37). Furthermore, another study showed that when trophoblasts come in contact with neutrophils, deactivation of neutrophils occurs (38). This suggests a defense mechanism of EVTs toward activated neutrophils, causing them to adopt an immunosuppressive phenotype.

Due to its highly positive charge, MPO can bind to extracellular GAGs. In the endothelial glycocalyx, MPO binds to the side chains of heparan sulfate, which causes the removal of syndecan-1 and disruption of glycocalyx formation (12). Other GAGs were found to affect MPO internalization into cells. It has been demonstrated that coating of the cell-derived extracellular matrix with sodium heparin results in higher MPO binding. Furthermore, when incubating MPO with sodium heparin, MPO bound to heparin in solution inhibited binding of MPO to the extracellular matrix (39). Based on these data, we selected sodium heparin as a blocker of MPO internalization and 4-ABAH as an irreversible inhibitor of MPO. We confirmed our hypothesis by demonstrating that the addition of heparin abolished MPO internalization into cells, whereas the addition of 4-ABAH did not affect internalization. Consistently, total MPO levels were depleted to that of untreated cells, indicating that heparin competitively binds MPO and completely eradicates its binding and internalization into cells. Finally, we showed that both heparin and 4-ABAH reduce MPO-mediated JEG-3 migration.

To our knowledge, this is the first study to examine the effect of MPO on JEG-3 cell migration in vitro. We found altered JEG-3 migration following MPO stimulation and internalization of MPO into cells. MPO localizes to the cytoplasm and nuclei of both tested cell lines, which were used as a model for EVT migration in vitro. The MPO effect was abolished by heparin and 4-ABAH. Overall, this study provides insight into mechanisms of the role of neutrophil-derived proteins that may lead to successful pregnancy. Furthermore, it suggests neutrophils and neutrophil-derived proteins, including MPO, as potent regulators of successful placentation. Therefore, MPO release, internalization, and function may represent potential therapeutic targets in pregnancy complications.

## CONCLUSIONS

To our knowledge, this is the first study to examine the effect of MPO on JEG-3 cell migration in vitro. We found increased JEG-3 migration following MPO stimulation and internalization of MPO into cells. MPO localized to the cytoplasm and nuclei in both tested cell lines, which were used as a model for studying EVT migration in vitro. The MPO effect was abolished by heparin and 4-ABAH pretreatment. Overall, this study provides insight into mechanisms, how extracellular MPO, predominantly derived from neutrophils, may alter EVT migration.

DB: Decidua basalis
ECIS: Electric cell-substrate impedance sensing
EMEM: Earle’s Minimum Essential Medium
EVT: Extravillous trophoblasts
FBS: Fetal bovine serum
ROS: Reactive oxygen species

## Author contributions

ZN.M designed and performed the experiments, analyzed and interpreted the results, crafted the figures, and wrote the manuscript. T.K. performed the experiments and analysis of the results. N.C-M. and P.V-C. assisted with the experiments. O.K. performed the Bioinformatics evaluation of the data. M.G and J.P. provided tissue blocks and JEG-3 cells. J.K. designed, supervised, and wrote the manuscript. All authors critically reviewed the manuscript and approved the submitted version.

## Funding

This work was supported by the FWF doctoral programs: DK-MOLIN (W1241), DP-iDP (DOC-31), and RespImmun (DOC-129) to PhD candidates ZN.M., N.C-M, and P.V-C. and OEAW (Doc Fellowship - 26477) to O.K., who were trained within the frame of the PhD Program in Molecular Medicine at the Medical University of Graz]. The work in the lab of J.K. was supported by the FWF [grant number P35294].

### Acknowledgements

We are grateful to Sabine Kern for her excellent technical assistance and to Marah Runtsch for proofreading the manuscript.

