## Supplementary Material for "Myeloperoxidase enhances the migration of human choriocarcinoma JEG-3 cells^1^"

Supplementary Figure 1

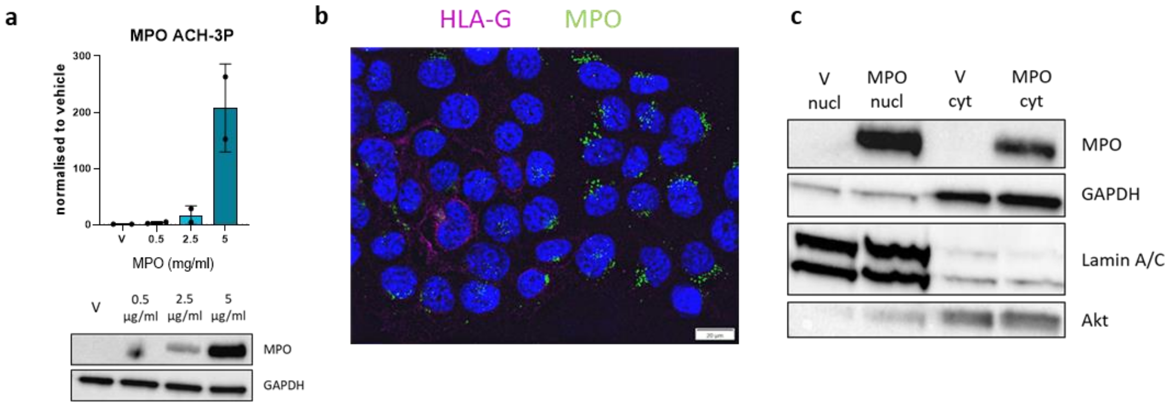


Supplementary Figure 1. MPO uptake and localization in ACH-3P cells

**a)** Western blot analysis of ACH-3P cells treated with various concentrations of MPO (0.5 µg/ml, 2,5 µg/ml and 5 µg/ml) presented as fold-change relative to vehicle (N=2). **b)** Representative immunofluorescence microscopy photos of 10 µg/ml MPO-treated cells. The cell nucleus was stained with DAPI and MPO is represented in green with AF488-labeled secondary antibody. HLA-G is represented in pink stained with AF647-labeled secondary antibody. **c)** Representative fractionation western blot images of MPO, GAPDH, Lamin A/C, and tAKT for vehicle and 5 µg/ml MPO-treated cells. Cytoplasmic and nuclear fractions were isolated. One-way ANOVA and Tukey's post hoc test were performed for multiple comparisons. Data are presented as the mean ± SD, *P < 0.05.
